# Learning from and improving upon high-throughput screens for protein fitness with Generative AI - Application to BBB-crossing AAV design

**DOI:** 10.1101/2024.12.06.627153

**Authors:** Ayan Kashyap, Kumari Soniya, Anubhooti, Subhalakshmi Kandavel, Kruthika Avadhani, Siva Kanishka, Manasvi Sharma, Prasad Chodavarapu

## Abstract

Deep Science is enabling high-throughput experimentation (HTE) to design novel biological entities with desired properties. E.g., blood-brain-barrier (BBB) crossing adeno-associated virus (AAV) vectors, needed for systemic delivery of gene therapies to brain cells, have been identified through innovative directed evolution assays such as M-CREATE and TRACER. But, even these high-throughput experiments are only able to explore a miniscule portion of the large design space of biological entities. In this paper, we introduce autograd based maximization of protein fitness (AutoMaxProFit) to learn from and improve upon protein designs generated with high-throughput screens. Using a transformer based generative AI network and protein language models, we improve upon the design of a variant previously discovered through HTE, to yield 2x better enrichment in brain endothelial cells, as estimated by molecular dynamics (MD) simulations. This shows that Deep Tech models can learn from the observations generated by Deep Science experiments and go on to find more optimal design candidates for application in Biopharma.

## Introduction

### Deep Science and Deep Tech are advancing Biopharma R&D

Every component of Biopharma R&D value chain - disease studies, target identification, elucidation and validation, to therapeutic design and optimization, synthesis, pharmaco-kinetic/ dynamic (PK/PD) and safety analysis - is undergoing rapid transformation, with the adoption of Deep Science and Deep Tech. Deep Science is allowing researchers to precisely perturb biological systems in high-throughput multiplexed protocols, and inspect/readout molecular and cellular states and activities. These advances on the science front are generating a wide-variety of large datasets that deep tech is helping analyze - to not only produce useful and novel insights, but also to integrate, model and generate novel designs. This process is visible across all disease areas as well as current and novel therapeutic modalities.

This paper fits into the same pattern, showing how Deep Tech can build on the datasets generated by Deep Science, for developing gene therapies that address genetic disorders impacting the brain.

### Gene therapies and viral vectors

Gene therapies aim to treat genetic disorders by delivering the missing functional genes into target cells. Adeno-associated virus (AAV) and lentivirus based vectors are commonly used in gene therapies as they have the natural ability to transduce cells and deliver genetic material safely.

A number of CNS disorders have genetic root causes. While local delivery of gene therapies into CNS is possible via intracranial and intrathecal administration into the brain and spinal cord respectively, systemic delivery via intravenous administration is more convenient for patients and care providers due to their non-invasive nature and are therefore, preferred over other delivery routes. However, intravenous administration requires high viral doses and are likely to provide relatively less efficient transduction of target cells, as crossing the blood-brain barrier (BBB) for reaching brain cells and blood-spinal cord barrier (BSCB) for reaching cells in the spinal cord, poses a major challenge for a successful gene therapy administration. Also, vectors used for systemic delivery should have minimal to nil penetration of cells other than the targeted cell types to minimize any off-target effects.

Success with intravenous administration of a gene therapy for spinal muscular atrophy (SMA), delivered using self-complementary (sc) AAV9 vectors [1] into mice and cats, led to human trials and eventual approval of the therapy. This encouraged research into similar vectors that can cross the BBB [2] for tackling genetic disorders of the brain such as neurodevelopmental and neurodegenerative disorders with genetic root causes.

### Directed evolution (DE) based innovations in BBB-crossing AAV

A number of studies over the years have focused on discovery of AAV variants that can cross the BBB and show tropism to specific cell types in the brain. (See [3] for a list of studies up to 2022). Studies such as CREATE [4], M-CREATE [5], TRACER [6] used directed evolution (DE) experiments in select mice models to identify specific 7-mer peptide inserts into the loop VIII (VR-VIII) region of AAV9 that bestow upon the resulting AAV9 variants, distinctly higher penetration into specific brain cell types when administered intravenously.

Subsequent studies (such as [7]) identified the specific receptors (such as LY6A in C57BL/6J mice) in brain endothelial cells that the BBB-penetrant AAV9 variants are binding to. This insight enabled the easier in vitro route to screen and discover more AAV variants that penetrate BBB in a broader set of mice models [8].

Combination of directed evolution, target receptor informed screening, and additional “semirational” capsid engineering informed by inspection of limited data available led to discovery of variants targeting BBB of other species such as non-human primates (NHP) [9, 10] and even humans ([11] with ex-vivo brain slices, [12] using transgenic mice).

Recently, in situ sequencing was used to create a spatial atlas of mouse CNS and the distribution of BBB-crossing AAV9 variants across different regions of the brain was studied [13], generating more data that can be mined with Deep Tech.

### Deep Tech enables further optimization

In all the discoveries described so far, while Deep Science enabled discovery of better variants, there was always a problem of limited data availability, in comparison to the size of the design space involved.

#### Problem of design space coverage

CREATE, M-CREATE and TRACER insert 7-mer peptides into the capsid sequence, looking for the best insert among 20^7^ (=1.28 billion) possible amino acid residue sequences or, equivalently, among approximately 3^7^*20^7^ (about 2.8 trillion) nucleotide sequences, by testing just 100 million nucleotide sequences. That represents a coverage of just 0.04% of the possible design space. Clearly, there’s an opportunity for improving upon the optimal sequences found through directed evolution, if only we can find a way to do so.

#### Generalization to protein fitness optimization

AAV optimization fits into a broader class of protein fitness optimization problems. Optimizing enzymes for catalytic activity as well as expression, stability, solubility and other properties required for developability is a common use case for protein fitness optimization methods.

#### Previous attempts

Previous attempts at application of AI/ML to AAV variant generation [14–16] focused mostly on generating a good dataset that’s likely to be representative of the fitness landscape over the design space. Predictive models developed on the experimental datasets were then used for screening [17–19] a large number of randomly generated sequences or sequences generated through saturation mutagenesis at select positions. Sequences with top predicted fitness scores were further validated with *in vitro* or *in vivo* studies.

#### Autograd based maximization of protein fitness (AutoMaxProFit)

In this paper, we describe a more direct approach, AutoMaxProFit, to finding optimal capsid sequences given the results of directed evolution experiments. Figure 1 summarizes the approach.

**Figure 1:**
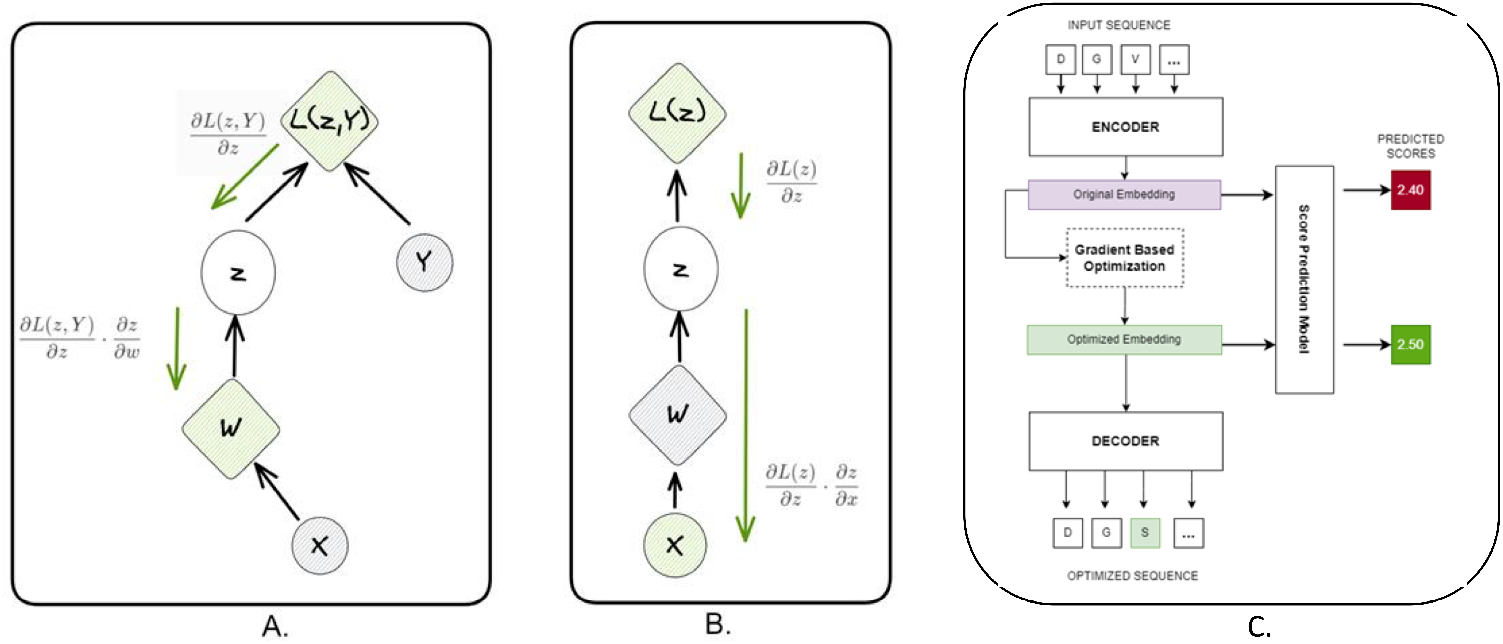
Schematics illustrating (A) Back-propagation of gradients during training of the model for prediction of protein fitness given sequence embedding computed by a protein language model. Loss function L compares prediction of fitness score z to ground truth Y. (B) Back-propagation of gradients during optimization. Loss function L is the predicted fitness score with its sign flipped to pose a maximization task as a minimization problem. (C) Process used to maxmimize fitness of a give protein in AutoMaxProFit.

AutoMaxProFit leverages the availability of pre-trained protein language models (pLMs) [20] to reduce the demands on training dataset size and coverage. A deep neural network based model is trained to predict protein fitness, given the representation of a protein sequence in the pLM embedding space.

To maximize protein fitness, AutoMaxProFit carries out gradient ascent in the embedding space, using autograd, the facility to compute gradients automatically in deep neural networks [21].

Finally, embeddings computed by the optimization process are mapped back into an amino acid sequence, using the de-masking trick all transformer based generative AI models use.

### MD for understanding tropism

We rely on Molecular dynamics (MD) simulations to validate the results from our optimization approach *in silico*.

MD simulations have proved to be useful in dissecting the molecular underpinnings of tropism, particularly by offering insights into how viruses interact with host cell receptors at an atomic level [22–25]. For instance, MD was successfully used in elucidating the role of the viral envelope protein gp120 in the binding of HIV-1 to CD4 receptor and co-receptors (CCR5 or CXCR4) on the host cells [26]. Static and dynamical analyses on MD trajectories were used to reveal how specific conformational changes, upon receptor binding, in gp120, a glycoprotein on the surface of HIV envelope, influence viral entry, shedding light on how alterations in the gp120 structure can impact tropism for different cell types [27].

Similar MD simulations have been performed on influenza virus to understand its tropism, e.g., how the hemagglutinin (HA) protein of influenza adapts to binding with sialic acid receptors on host cells. By simulating different HA variants, researchers could identify structural features critical for binding to specific sialic acid linkages, which correlate with the virus’s ability to infect various host species [25]. Recently, MD studies have been extensively used in understanding SARS-CoV2 binding with ACE2 receptors and the effect of different mutations on binding [23]. These examples highlight how MD simulations can reveal the fine details of virus-receptor interactions and provide a platform for exploring how mutations or structural changes influence tropism.

In the present work, conventional MD simulations (cMD) were employed to quantify the binding and stability of AAV9 vectors when complexed with LY6A receptors on endothelial cells. While RMSD was used to account for the stability of the AAV9-LY6A complex, binding free energies and buried surface area were used to quantify LY6A binding relative to the known AAV9 capsids exhibiting tropism. Results are discussed in later section.

## Results

### Performance of predictive model

#### Learning from directed evolution

Results from M-CREATE are available in the form of enrichment (ER) scores, representing the relative success of a particular capsid sequence in penetration of a target tissue/cell type. We trained a model to predict the enrichment score of a capsid for endothelial cells in the brain, given its amino acid sequence.

To featurize the sequence, we relied on embeddings generated by ProtT5 [20], a protein language model that was trained on more than 2 million protein sequences, using an encoderdecoder transformer architecture [28].

We reduced the dimensionality of the capsid sequence embedding from 525 (residues in capsid sequence) * 1024 (dimensions/residue in ProtT5 embeddings) to 30 * 1024 by retaining only 30 residues in loops IV and VIII (including those in the 7-mer insert) that showed significant variation of embedding within the training set.

We used a 1-D convolutional neural network (CNN) shown in Figure 7, inspired by the best performing architecture used for secondary structure prediction in ProtT5.

Figure 2 shows (a) the distribution of ER scores for endothelial cells in M-CREATE dataset and (b) the performance of the ER score prediction model. Top-k recall values are 1, 0.6 and 0.56 for k=1, 10 and 50 respectively.

**Figure 2:**
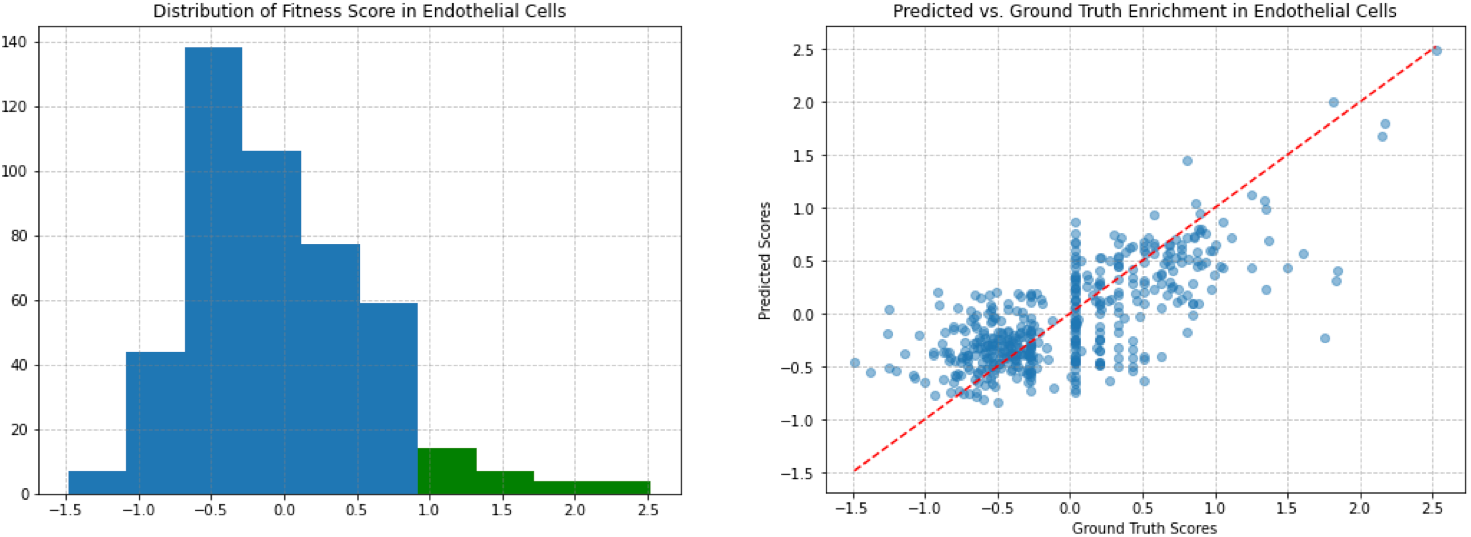
Distribution of ground truth enrichment scores and Ground truth vs prediction scatter plot for endothelial cells

### Capsid optimization and validation

#### Improving upon directed evolution

To improve upon the best sequences found by M-CREATE, we conducted a numerical optimization process in the (dimensionally reduced) sequence embedding space to find the embeddings that produced the highest enrichment score prediction (using the model described in previous section) and subsequently, decoded the embeddings to produce the optimal residue sequence.

For numerical optimization, we used back-propagation with gradient descent enabled by automatic differentiation in PyTorch [21], just as in training of the predictive model, except that the gradients computed during optimization were with respect to the 30 * 1024 input embeddings, and not wrt. the parameters (weights and biases) of the 1-D CNN.

Loss function for the optimizer combined predicted enrichment score (with its sign flipped to pose a maximization task as a minimization problem) along with a penalty for straying outside the embeddings sub-space covered by the training dataset, as shown in Figure 8.

Figure 3 shows how we decoded the embeddings found by the optimization process to yield the (predicted) best enrichment score. Post decoding, we checked to make sure residues outside the 7-mer insert stayed identical to the residues in the wildtype sequence. Any embeddings that did not decode well under this constraint were discarded in favour of next best scoring embeddings that did satisfy the constraint.

**Figure 3:**
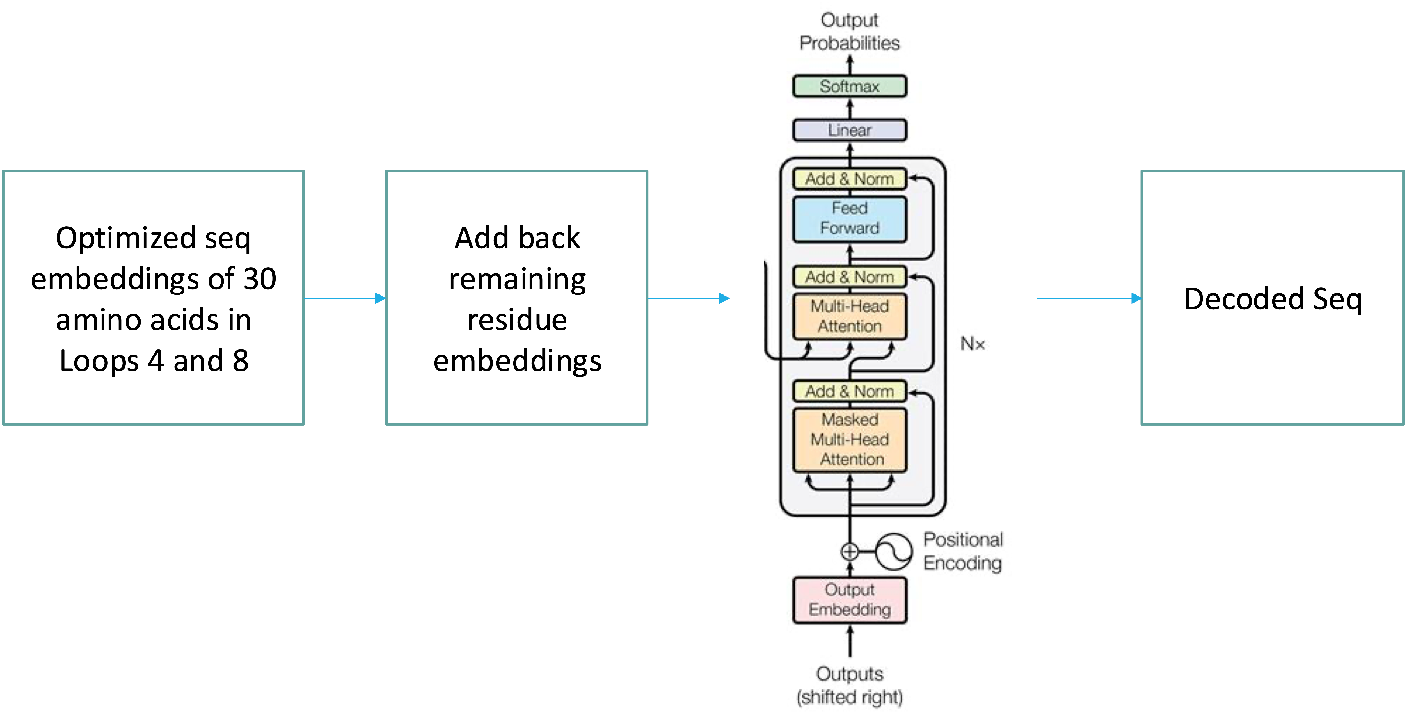
Decoding the sequence embeddings found by the optimization process to yield the highest enrichment score prediction

Table 1 shows the changes optimizer made to maximize the enrichment score for brain endothelial cells, starting from TLQLPFK, a top 7-mer insert found by M-CREATE to yield a good enrichment score for multiple brain cell types.

**Table 1:** 7-mer sequences for AAV9 loop VIII insert generated by AutoMaxProFit, starting from TLQLPFK, a top insert found by M-CREATE. Also shown are the enrichment scores predicted.

| 7-mer | Predicted Enrichment Score |
| --- | --- |
| TLQLPFK | 2.81 |
| SLQAPFK | 3.04 |
| SLQAPWK | 2.87 |
| SLQAQQQ | 3.20 |
| SLQALQA | 3.39 |

#### Validation of optimized capsid fitness through MD simulations

Trimeric engineered AAV9 capsid protein was complexed with matured LY6A receptor using Haddock3. Haddock3 can be accessed through this repository https://github.com/haddocking/haddock3. Selected docked complex structures were subjected to 100 ns cMD simulations.

Figure 4 shows the trimeric AAV9 variant structure. The 7-mer peptide inserts were placed in loop VIII of the protein. In the trimeric structure, loop IV, loop V and loop VIII extend outwards to interact with receptors. The trimeric AAV9 protein is bundled together by intertwining the loop V from the adjacent chain with loop IV and loop VIII of the first chain.

**Figure 4:**
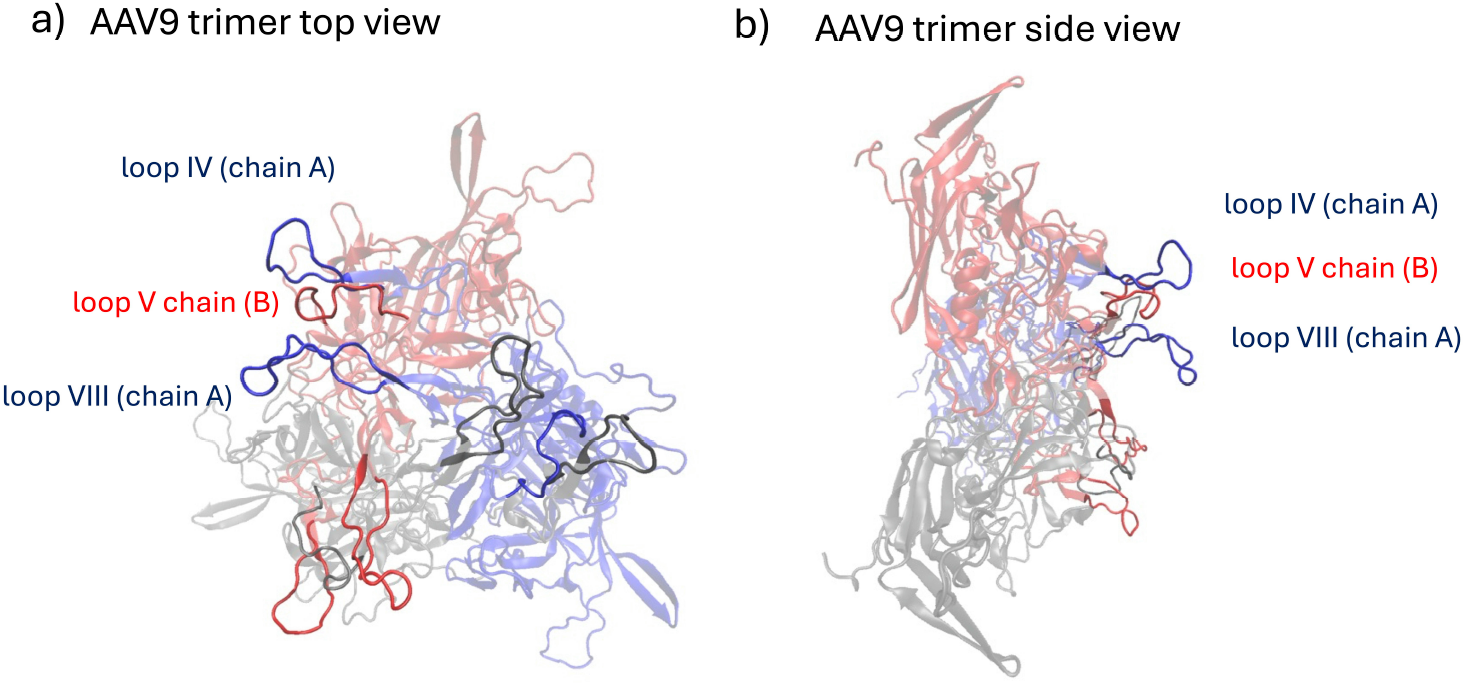
Positions of loops IV, V and VIII in the trimeric AAV9 as seen in a) top view and b) side view

To assess the stability of the complex, we monitored fluctuations in the root-mean-square deviation (RMSD) of the complex’s structure over the length of the simulation, shown in Figure 9. We used binding free energy and buried surface area (BSA) as quantitative measures to evaluate the optimised capsid. For comparison, binding free energies and BSA values for known penetrants and non-penetrants, obtained from the M-CREATE and TRACER platforms, were used as controls.

Table 2 shows the quantitative results obtained from analysis of MD trajectories. The capsid optimized by AutoMaxProFit showed a 2x improvement in binding strength.

**Table 2:** Comparison of relative binding free energy and BSA of in-house optimised capsid to the known controls from M-CREATE and TRACER.

| 7-mer system | Binding free energy (k-cal/mol) | BSA ($\text{\AA}^2$) |
| --- | --- | --- |
| M-CREATE penetrant | -41 | 1231.2 |
| M-CREATE non-penetrant | -30 | 626.2 |
| 9P16 (TRACER penetrant) | -49 | 776.2 |
| 9P09 (TRACER non-penetrant) | -36 | 555.7 |
| SLQALQA (AutoMaxProFit designed capsid) | -86 | 1420.8 |

A greater binding free energy and larger buried surface area as compared to the known penetrants/controls further help in validating the prediction made by AutoMaxProFit. Representative frames from a 100 ns MD trajectory are depicted in Figure 5. The left and middle images are the BBB penetrating controls, while the rightmost image illustrates the optimised sequence. The lower panel displays known BBB non-penetranting capsids.

**Figure 5:**
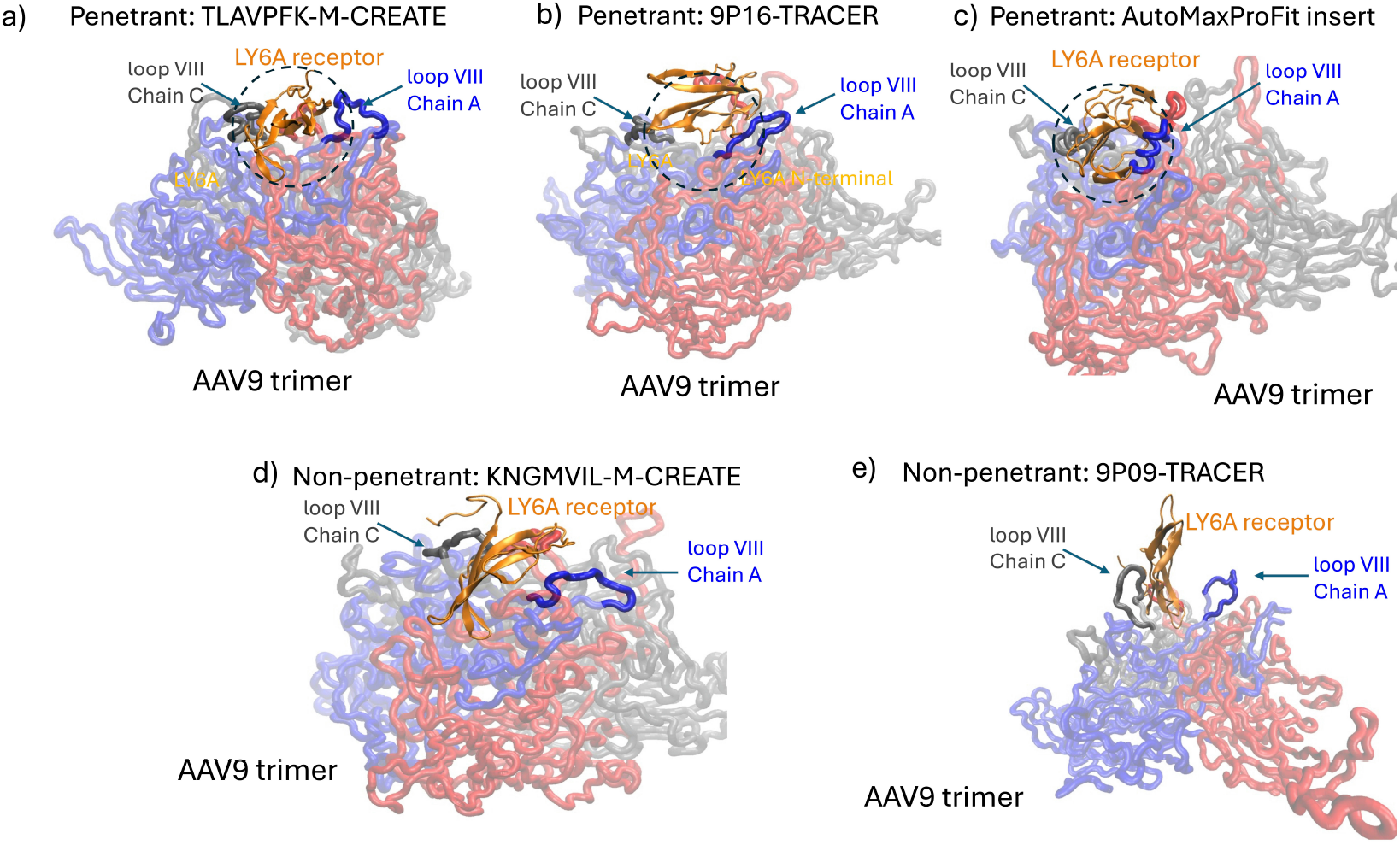
Final frames from a 20 ns MD trajectory. a) Optimal penetrant from M-CREATE complexed to LY6A; b) Optimal penetrant from TRACER complexed to LY6A; c) Optimal penetrant from AutoMaxProFit complexed to LY6A; d) A non-penetrant from M-CREATE complexed to LY6A; and e) A non-penetrant from TRACER complexed to LY6A. Notably, penetrants exhibit extensive engagement with the 7-mer inserts resulting in improved binding free energy and a larger buried surface area upon complex formation. In contrast, non-penetrants show weaker interaction with loop VIII inserts in the trimer.

## Discussion

In this paper, we illustrated how Deep Tech can build on the discoveries made by directed evolution studies in vivo. Similar value can be added on top of discoveries made in other studies that leveraged knowledge of BBB receptors and spatial omics studies that uncovered tissue distributions of penetrating AAV variants.

The usefulness of AutoMaxProFit approach extends much beyond design of BBB-crossing AAV variants. As noted earlier, enzyme optimization for maximizing catalytic activity while ensuring other properties are within desired ranges, can also be accomplished similarly.

In this study, we have limited validation to a rather simplistic MD simulation based approach. Multiple replicas and better sampling are needed to obtain more rigorous inferences in MD.

Given the promise of the results from this study, we hope to repeat the same on AAV variants identified via high-throughput experimentation, for penetration of BBB in NHP and humans. We also plan to conduct to a similar proof-of-concept study for enzyme optimization.

## Methods

### Featurization of the AAV capsid sequence

We scaled each dimension of the embeddings to [0,1] using minimum and maximum values found for each dimension across all the training dataset.

**Figure 6:**
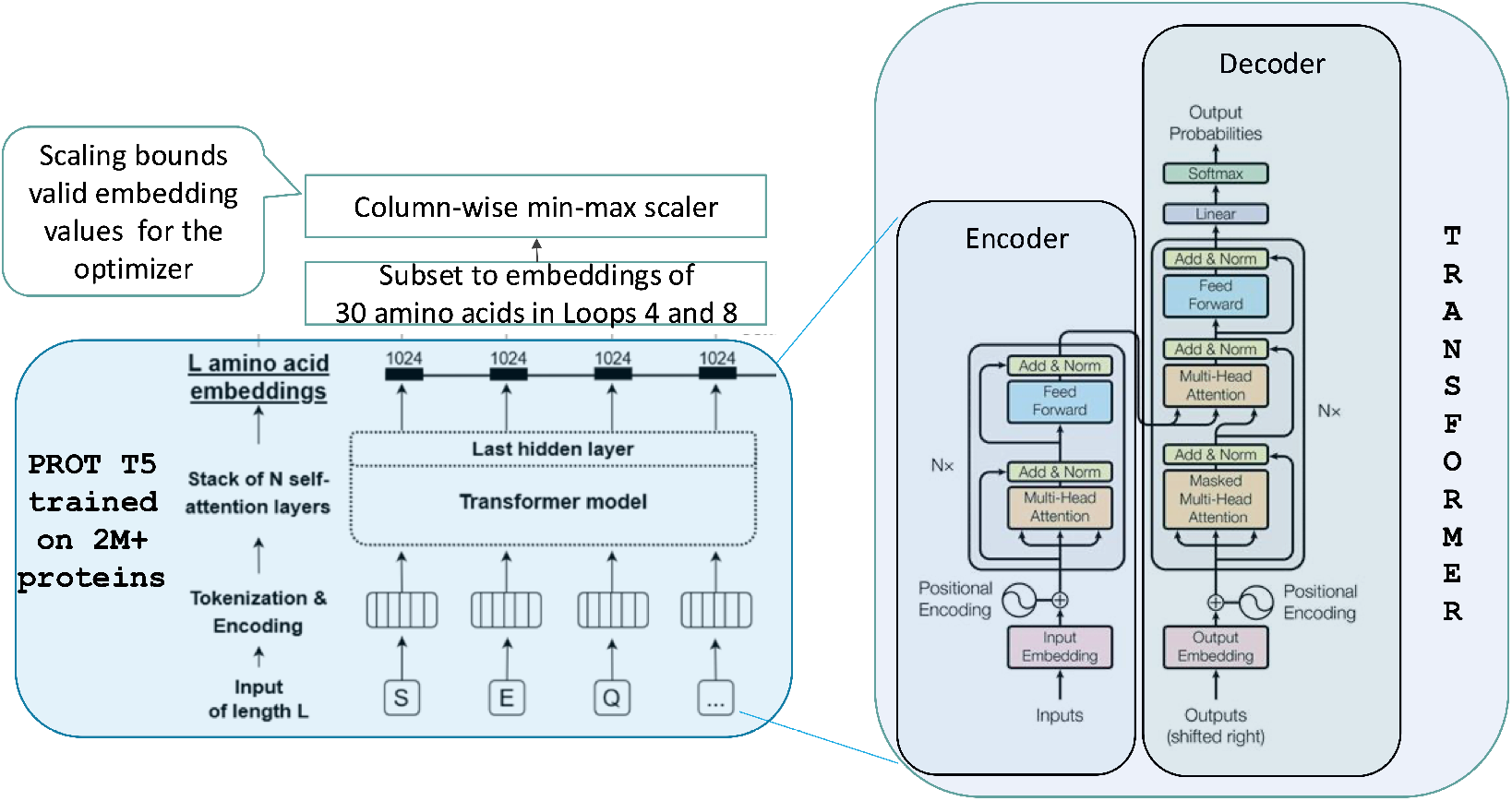
Featurization of AAV capsid sequence using a protein language model

### Prediction model for enrichment score

**Figure 7:**
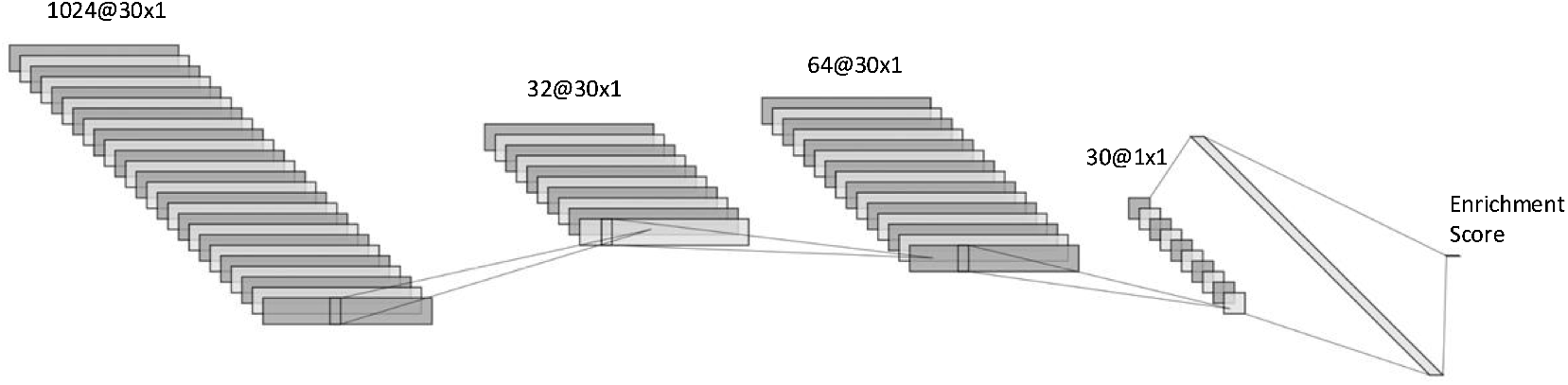
1-D convolutional neural network (CNN) architecture used for training models to predict enrichment score of a given capsid sequence, for a target tissue/cell type.

### Optimization using prediction model

**Figure 8:**
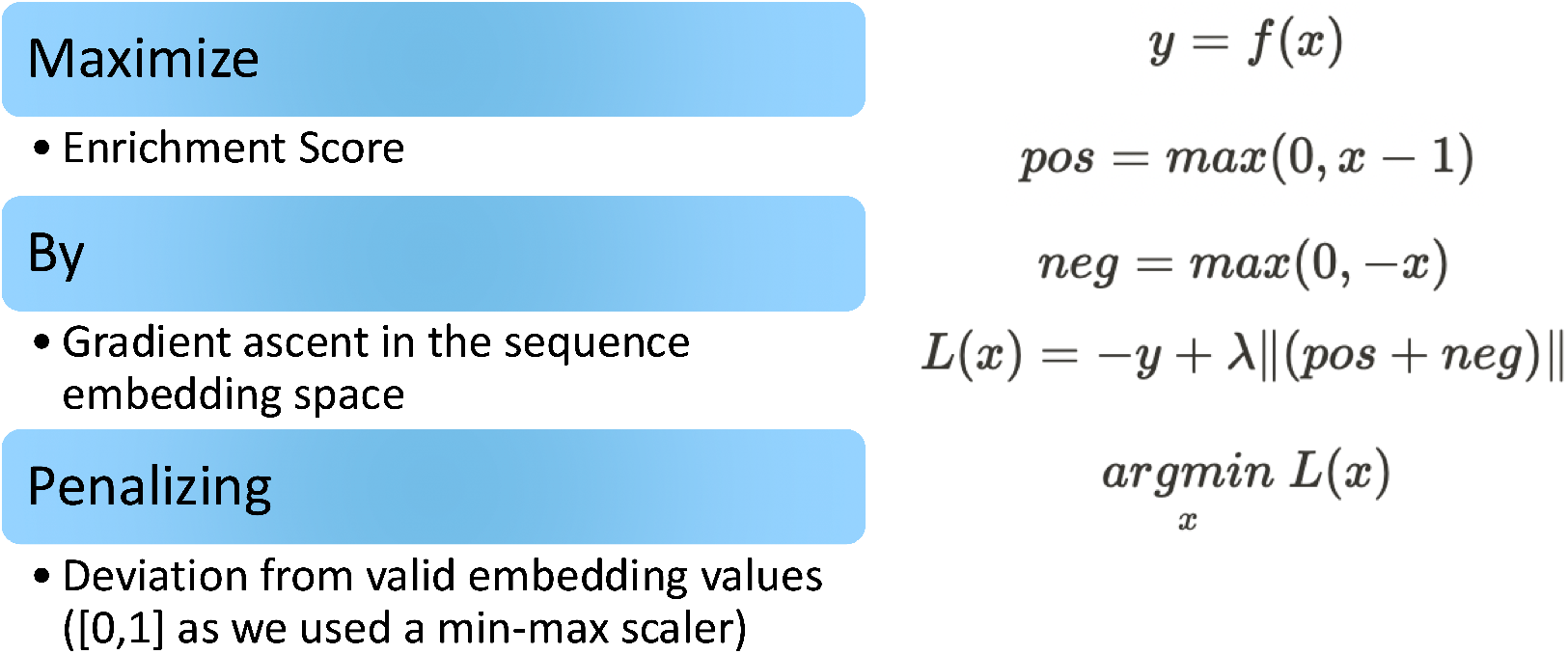
Loss function used in optimization

### Molecular dynamics (MD) simulations

#### Structure preparation

The structure for PHP.eB variant of the AAV9 vector was extracted from PDB ID 7WQO [29]. To study tropism, the essential structure is the trimeric AAV9 in complex with the receptor LY6A. Experiments using AF2-multimer [30] (via Colabfold [31]) showed that while it can reliably predict a monomeric AAV9-LY6A complex, it struggles with predicting a complex involving three AAV9 and one LY6A chains. To address this, we constructed a trimeric AAV9 model using VIPERdb [32]. This trimeric AAV9 protein was complexed with LY6A using HADDOCK3.

#### Docking

Rigid-body docking was performed using 1000 initial models, incorporating ambiguous and unambiguous restraints to guide interactions between the LY6A receptor and the trimeric AAV9 capsid protein.

After rigid-body docking, clusters were ranked using FCC-based clustering, and the top 10 models were selected for further refinement in the flexible refinement stage, where backbone flexibility was introduced with ssdihed = “alphabeta” and an epsilon (dielectric constant) of 2.0. A 10% failure tolerance was permitted during flexible refinement.

Final models underwent molecular dynamics refinement, further refining loop flexibility and intermolecular contacts while maintaining predefined restraints to ensure structural fidelity.

Based on receptor interactions with the desired loop 8, the best-ranked starting structures were chosen for molecular dynamics simulations.

Above protocols were used to prepare five systems, including the four that served as the controls for tropism analysis via molecular dynamics (both penetrants and non-penetrants of the blood-brain barrier) and one that was AutoMaxProFit optimised penetrant for endothelial cells.

#### MD Simulation Details

The molecular dynamics (MD) simulations of the trimeric AAV9LY6A complex in water was carried out in GROMACS 2022.3 [33, 34].

For protein, amber.ff99SB [35] force field was used. The complex was solvated in a dodecahedron box containing SPC/E water model under neutral conditions. The prepared systems contain more than 300,000 atoms in each case.

To ensure accurate dynamics, the system was first energy minimised using the steepest descents algorithm, followed by equilibration in two phases: NVT ( constant volume and tempearature) and NPT (constant pressure and temperature). During NVT equilibration, temperature was controlled using the velocity rescaling thermostat [36] at 300 K. In the subsequent NPT equilibration, pressure was maintained at 1 bar using the Parrinello-Rehman barostat [37].

Production simulations were carried out in the NPT ensemble with a time step of 2 fs.

The Particle Mesh Ewald (PME) algorithm [38] was used to account for the long-range electrostatic interactions. Hydrogen atoms were constrained using the LINCS algorithm [39]. MMGBSA and BSA calculations were done on the generated trajectories to obtain the trimeric AAV9-LY6A binding affinities and the resulting surface area due to complexation. gmx_MMPBSA tool [40] was used for binding free energy calculations. Dr_SASA [41] was used on the final frame from MD trajectory to evaluate buried surface area between the trimeric AAV9 and the LY6A receptor. Root mean square deviations were calculated on the MD trajectory using GROMACS.

**Figure 9:**
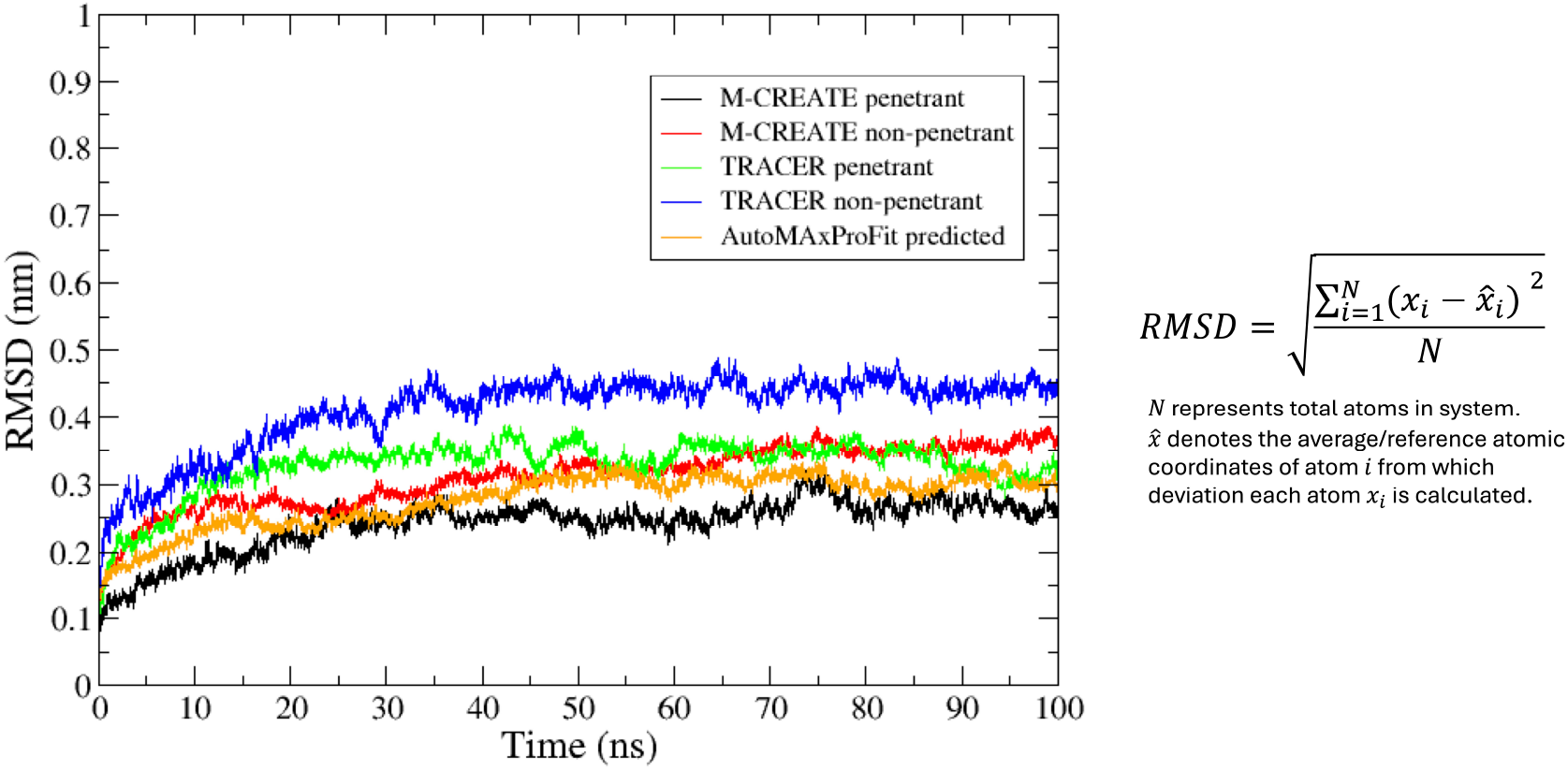
RMSD fluctuations for trimeric AAV9-LY6A complexes with time

